# The mTORC1/S6K/PDCD4/eIF4A axis determines outcome of mitosis

**DOI:** 10.1101/794545

**Authors:** Mohamed Moustafa-Kamal, Thomas Kucharski, Wissal El Assad, Valentina Gandin, Yazan Abas, Bhushan Nagar, Jerry Pelletier, Ivan Topisirovic, Jose G. Teodoro

**Affiliations:** Goodman Cancer Research Center, McGill University, Montréal, Québec, Canada; Department of Biochemistry, McGill University, Montréal, Québec, Canada; Department of Microbiology and Immunology, Montréal, Québec, Canada; Lady Davis Institute for Medical Research, Sir Mortimer B. Davis-Jewish General Hospital, and Department of Oncology, McGill University, Montréal, Québec, Canada

**Keywords:** mTORC1, Raptor, cell cycle, mitosis, PDCD4, eIF4A

## Abstract

mTOR is a serine/threonine kinase which acts a master regulator of cell growth and proliferation. Raptor, a scaffolding protein that recruits substrates to mTOR complex 1 (mTORC1), is known to be phosphorylated during mitosis, but the significance of this phosphorylation remains largely unknown. Here we show that raptor expression and mTORC1 activity are dramatically reduced in mitotic arrested cells across a variety of cancer and normal cell lines. Prevention of raptor phosphorylation during mitosis resulted in reactivation of mTORC1 in a rapamycin-sensitive manner. Importantly, expression of a non-phosphorylatable raptor mutant caused a dramatic reduction in cytotoxicity of the spindle poison Taxol. This effect was mediated via degradation of Programmed Cell Death Protein 4 (PDCD4), a tumor suppressor protein that inhibits eIF4A activity and is negatively regulated by the mTORC1/S6K pathway. Moreover, pharmacological inhibition of eIF4A was able to enhance the effects of taxol and restore sensitivity in Taxol resistant cancer cells. These findings indicate that the mTORC1/S6K/PDCD4/eIF4A axis has a pivotal role in death vs. slippage decision during prolonged mitotic arrest and may be exploited to gain a clinical benefit in treating cancers resistant to anti-mitotic drugs.

## Introduction

In order to maintain proper tissue homeostasis, cells need to coordinate both growth (increase in cell mass) and proliferation (increase in cell number). The two processes are linked via the evolutionarily conserved TOR (Target Of Rapamycin) signaling pathway, which integrates a variety of extracellular signals and intracellular cues including hormones, growth factors and nutrients to coordinate growth and proliferation with metabolic activity in the cell. In mammals mechanistic/mammalian TOR (mTOR) nucleates two different large signaling complexes: mTOR complex 1 (mTORC1) and 2 (mTORC2). mTORC1 consists of mTOR, raptor (regulatory associated protein of mTOR), mLST8 (mammalian lethal with sec-13), PRAS40 (proline-rich AKT substrate 40 kDa), and DEPTOR (DEP domain-containing mTOR interacting protein). mTORC1 stimulates anabolic processes such as protein synthesis and energy production. mTORC2 is composed of mTOR, rictor (raptor independent companion of mTOR), mLST8, mSIN1 observed with rictor-1) and controls cytoskeletal organization and cell survival (Laplante and Sabatini, 2012; Mossmann et al., 2018; Saxton and Sabatini, 2017).

In yeast, TOR primarily regulates cell growth and secondarily impacts on proliferation (Barbet et al., 1996; Conlon and Raff, 2003). In mammals, mTORC1 impacts on both cell growth and proliferation, which is mediated by the eukaryotic translation initiation factor 4E (eIF4E)-binding proteins (4E-BPs) and ribosomal protein S6 kinases (S6Ks), respectively (Dowling et al., 2010). 4E-BPs and S6Ks mediate the effects of mTORC1 on protein synthesis. During cap-dependent translation initiation, mRNA is recruited to the ribosome via the eIF4F complex, which comprises a cap-binding subunit eIF4E, large scaffolding protein eIF4G and DEAD box RNA helicase eIF4A, which facilitates scanning of the ribosome for the initiation codon. Phosphorylation of 4E-BPs by mTORC1 stimulates their release from eIF4E, which allows eIF4E-eIF4G association and the assembly of the eIF4F complex, thereby increasing translation initiation rates (Roux and Topisirovic, 2018; Sonenberg and Hinnebusch, 2009). S6Ks phosphorylate a number of components of the translational machinery and related regulators such as ribosomal protein S6, eIF4B, eEF2K and PDCD4, an inhibitor of eIF4A (Roux and Topisirovic, 2018; Zoncu et al., 2011).

Previous studies indicated that raptor has a role in mediating mTORC1 assembly, recruiting substrates, and regulating mTORC1 activity (Hara et al., 2002; Yip et al., 2010). Recent studies have demonstrated the importance of phosphorylation of raptor on various sites in the regulation of mTOR signaling by pro- and anti-proliferative signals. Phosphorylation by Rsk at S721 (Carriere et al., 2008) as well as by mTOR at S863 (Foster et al., 2010) have been shown to enhance mTORC1 activity, whereas phosphorylation at S722 and S792 by AMPK create 14-3-3 binding sites and suppress mTORC1 activity (Gwinn et al., 2008). Raptor has also been shown to be heavily phosphorylated in mitosis on at least 9 conserved sites down stream of cyclin-dependent kinase 1 (cdk1) and glycogen synthase kinase 3 (GSK3) (Gwinn et al., 2010; Ramirez-Valle et al., 2010). These reports showed that mTORC1 activity is needed for mitotic progression despite the reportedly decreased mitotic activity of two of the upstream activators of the mTORC1 pathway, AKT and MAPK pathways (Alvarez et al., 2001; Ramirez-Valle et al., 2010). Notwithstanding these findings, the significance of mitotic phosphorylation of raptor and the role of mTORC1 in mitotic progression remains poorly understood.

In the current study we observed, somewhat unexpectedly, that mTORC1 activity in mitosis is dramatically reduced. Furthermore, we show that multisite mitotic phosphorylation of raptor leads to a reduction in mTORC1 activity. A nonphosphorylatable raptor mutant reactivates the mTORC1 complex and promotes extended survival of cells challenged with taxol. Finally, we demonstrate that mTORC1 delays cell death under mitotic arrest by inducing degradation of PDCD4 pro-apoptotic protein and subsequently bolstering eIF4A activity. These results highlight a previously unappreciated role of the mTORC1/S6K/PDCD4/eIF4A axis in mitosis and suggest that targeting this axis may increase anti-neoplastic efficacy of mitotic poisons.

## Results

### mTORC1 activity is decreased during mitotic arrest

In order to examine the activity of mTORC1 complex in the context of prolonged mitosis (mitotic arrest), HeLa cells were synchronized using thymidine followed by release into nocodazole (Noc) and analyzed by immunoblotting at indicated time points post release (Figure 1A). This experiment revealed that the cells that progress into mitosis, as evidenced by the appearance of phosphorylated cdc27 and geminin, gradually decrease S6K phosphorylation at Thr-389, which is a well-established mTORC1-specific phosphorylation site (Figure 1A). Concomitantly, we observed an upward electrophoretic mobility shift and a decrease in raptor levels (Figure 1A, lanes 6-8). Treatment of cell extracts with λ phosphatase reversed the upward mobility shift of mitotic raptor confirming that it is caused by phosphorylation (Supp. Fig. 1A). In contrast to S6K, phosphorylation of 4E-BP1 was increased in mitosis, which is consistent with a previous report showing that CDK1, and not mTORC1, phosphorylates 4E-BP1 in mitosis (Velasquez et al., 2016). Accordingly, active-site mTOR inhibitor, torin 1, inhibited the phosphorylation of S6K both in mitosis and interphase and attenuated 4E-BP1 phosphorylation in interphase but not in mitosis (Figure 1B).

**Figure 1:**
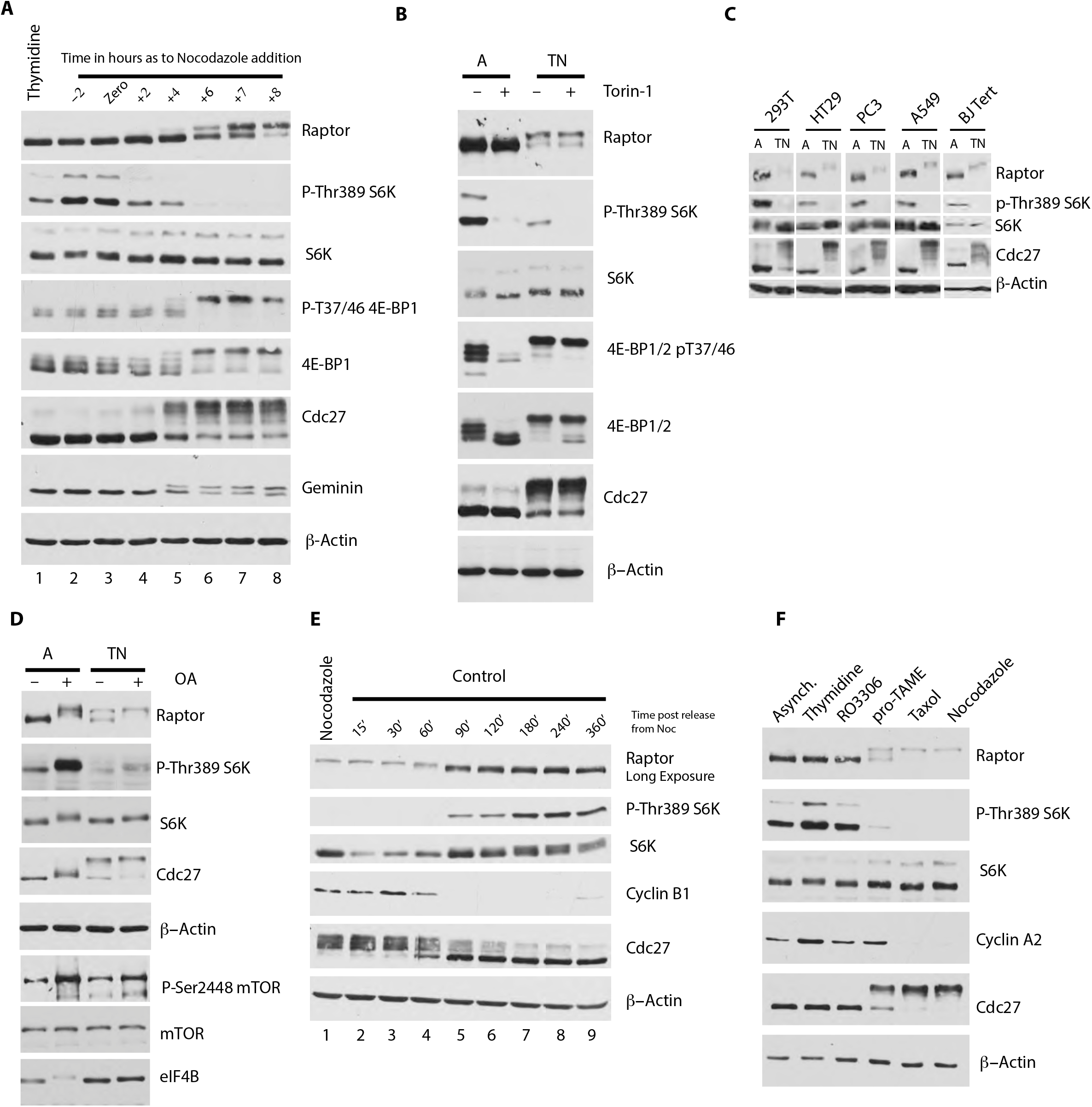
**A.** Immunoblot analysis of HeLa cells synchronized using Thymidine followed by release into Nocodazole. Whole cell extracts were analyzed for expression of raptor and signalling pathway targets downstream mTORC1, like phospho-Thr389-S6K and phospho-4E-BP1, following release from Thymidine into Nocodazole at the indicated times. Anti-Cdc27 and anti-Geminin are included as a marker for onset of mitosis and β-Actin as a loading control. **B.** Immunoblot analysis of HeLa cells that were either asynchronous (A) or synchronized using Thymindine/Nocodazole protocol (TN). Torin 1 (200nM) were added for 3hours before harvesting. **C.** Immunoblot analysis of different cell lines that were either left unsynchronized (A) or synchronized with Thymidine/Nocodazole protocol (TN). Floating mitotic cells were collected by shake-off. **D.** Immunoblot of HeLa cells synchronized as in **B**. Two hours before harvesting cells were treated with Okadic acid (OA). Phospho-Ser2448 mTOR and eIF4B were used as controls for the phosphatase treatment. **E.** Immunoblot analysis of HeLa cells synchronized using Thymidine-Nocodazole protocol then Nocodazole was washed of mitotic cells and released into fresh media and followed at the indicated times. Cyclin B1 was used as a marker for mitotic progression and exit. **F.** HeLa cells were synchronized with Thymidine for 20 hours, and then released into media with RO3306 (CDK1 inhibitor), pro-TAME (an APC/C inhibitor), Taxol, and Nocodazole for 12 hours. Anti-Cyclin A2 was used as a control for pro-TAME.

These data suggest that in contrast to previously published findings (Gwinn et al., 2010; Ramirez-Valle et al., 2010), mTORC1 activity appears to be downregulated during mitosis or mitotic arrest. Notably, previous studies used HaCat cells, which are primary human fibroblast that are technically harder to synchronize in mitosis than cancer cell lines employed in the present study (Ramirez-Valle et al., 2010). HaCat cells required a mitotic shake off protocol to prevent contamination with non-mitotic cells and we speculate that this may explain discrepancy of our results with the previous report (Suppl. Fig. 1C) (Ramirez-Valle et al., 2010). Importantly, we detected comparable downregulation of mTORC1 activity as illustrated by a decrease in S6K phosphorylation in mitosis in all cell lines tested including HEK293T, PC3, HT29, A549, and telomerase-immortalized BJ-tert human fibroblasts (Figure 1C). This was paralleled by electromobility shifts and decrease in raptor levels. Moreover, mTORC1 activity was also reduced during mitosis in TSC2 null MEFs, which uncouples mTORC1 from the canonical upstream pathways that activate/deactivate it during interphase. (Suppl. Fig. 1C).

The PP2A phosphatase has been reported to dephosphorylate mTORC1 substrates (Apostolidis et al., 2016). The PP2A inhibitor, okadaic acid, however, did not rescue S6K phosphorylation in mitosis, as compared to interphase cells (Figure 1D), thus excluding the possibility that the observed decrease in S6K phosphorylation was caused by increased PPA2 activity in mitosis. We also excluded the possibility that the changes in S6K phosphorylation stem from inadvertent effects of extended mitotic arrest caused by Noc treatment. To this end, HeLa cells that were released from mitotic arrest into fresh media and collected over time as they exit mitosis into G1-phase (Figure 1E). Strikingly, S6K phosphorylation at Thr-389 started to increase only after cells completely exited mitosis as indicated by loss of cyclin B levels and Cdc27 phosphorylation (Fig. 1E, lane 5-9), concomitant with loss of electrophoretic mobility shift of Raptor. Finally, we used the same protocol to collect mitotic HeLa cells after treatment with other agents that cause prolonged mitotic arrest including the APC/C inhibitor, pro-TAME and Taxol, which produced shifts in electrophoretic mobility of raptor and decrease in S6K phosphorylation that were comparable to those observed using Noc (Figure 1F). Taken together, these results show that mTORC1-dependent phosphorylation of S6K is diminished during mitotic arrest, which correlates with phosphorylation of raptor.

### Raptor protein is downregulated during mitotic arrest

To further characterize the regulation of raptor during mitosis, we set out to determine whether the mitotic loss of mTORC1 activity occurs during normal mitosis or if it is due to prolonged mitotic arrest. To address this question, HeLa cells were arrested at G2/M using the cdk1 inhibitor, RO3306, and released into fresh media containing either Noc or vehicle (DMSO). These experiments revealed that under conditions of normal mitosis, raptor is phosphorylated as efficiently as in the presence of Noc (Figure 2A, compare lanes 4 and 5). However, the mitotic decrease in raptor levels occurred only in under conditions of sustained mitotic arrest induced by Noc (for example, compare lanes 6 and 7). The observation that raptor levels are decreased in a manner that is coupled with phosphorylation suggest that the protein is actively degraded in a cell cycle dependent manner. We therefore investigated whether raptor is degraded by the proteasome. Treatment with the proteasome inhibitor, MG132, did not to rescue mitotic levels of raptor, but did rescue other proteasome targets including wee-1 and Cyclin A2 (Figure 2B, compare lanes 5 and 6). Other possible mechanisms of raptor degradation including caspase cleavage or lysosomal degradation were tested as causes of reduced mitotic raptor levels. As with MG132, pan-caspase (zVad-FMK) or lysosome activity inhibitors (chloroquine, Bafilomycin A, or NH_2_Cl) also failed to restore mitotic raptor levels (Fig. 2C and 2D). The possibility that we failed to detect raptor due of technical artifacts caused by epitope masking through phosphorylation or protein insolubility were also ruled out as neither lysing mitotic HeLa cells in Laemmli buffer nor treatment of lysates with λ-phosphatase could rescue the raptor levels (Supplemental Fig. 2A, 1A). We then assessed the level of raptor mRNA using RT-PCR which showed a modest decrease (approximately 25%) as cells progress in mitosis which is insufficient to account for the decrease in raptor protein levels (Supplemental Fig. 2B).

**Figure 2:**
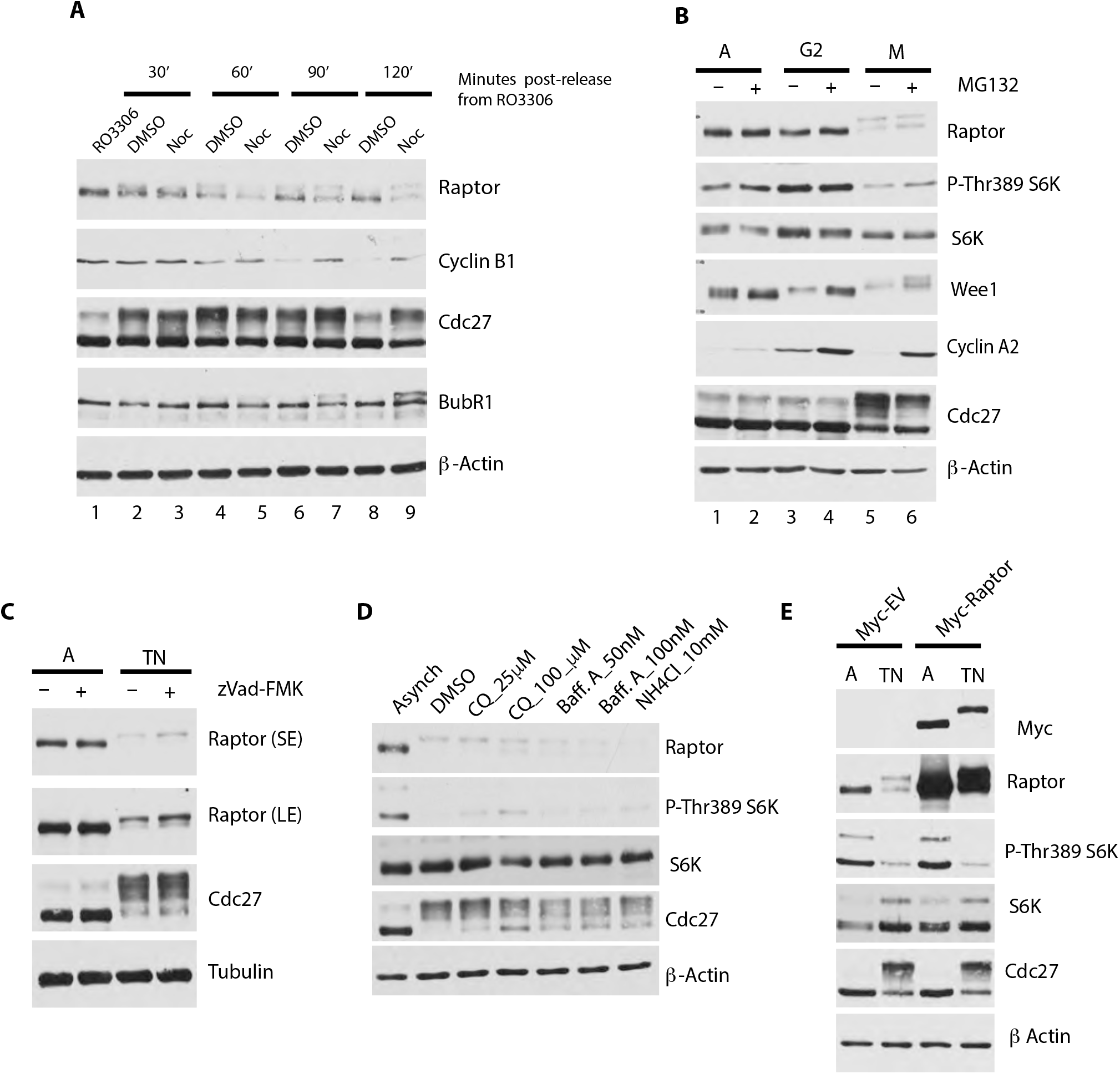
**A.** Immunoblot analysis of HeLa cells synchronized with the cdk1 inhibitor RO3306 and released for the indicated length of time in vehicle control (DMSO) or in Nocodazole. Cyclin B1 was used as a marker for mitotic exit. Phosphorylated Cdc27 and BubR1 were used as markers for mitosis. **B.** HeLa cells were either left untreated (A) or treated with RO3306 for 19 hours (G2) and then released into fresh media containg Nocodazole to trap them in mitosis (M). One set of all plates was treated with MG132 for 3hours. immunoblot analysis for expression of Wee 1 and cyclin A2 known to be degraded during mitosis were used as controls. **C.** Immunoblot analysis of asynchronous (A) or Thym/Noc (TN) synchronised HeLa then treated with vehicle control or the pan-caspase inhibitor zVad-FMK for the last 12 hours of treatment. D. HeLa cells were treated as in C, but then released into fresh media containg Nocodazole plus vehicle control or three different lysosome inhibitors: cloroquine (CQ), Baffylomycin A (Baff A), and ammonium chloride (NH_4_Cl). **E-**HeLa cells were transfected with pRK5-myc empty vector or Raptor WT and synchronized as in C.

The above data show that there is a near complete loss of mTORC1 activity at mitosis paralleled with simultaneous phosphorylation and reduction in raptor protein levels. We therefore determined whether expression of exogenous raptor cDNA driven by a CMV promoter could rescue mTORC1 activity in mitosis. Although transfection of raptor cDNA was able to rescue levels of mitotic raptor, it failed to rescue the mTORC1 activity as evidenced by the continued absence of Thr-389 S6K signal (Figure 2E). Moreover, transfected myc-tagged raptor appeared to be almost entirely in the slower migrating, phosphorylated form. Taken together these data show that mTORC1 activity is inhibited at mitosis through a phosphorylation dependent mechanism. Furthermore, conditions of prolonged mitotic arrest appear to negatively regulate levels of raptor protein through an unknown mechanism. Ectopic expression of raptor could not rescue mTORC1 activity suggesting that phosphorylation of raptor is sufficient to inhibit mTORC1 activity during mitosis.

### Raptor phosphorylation regulates mTORC1 dimerization

mTORC1 is thought to act as an obligate dimer with raptor having a crucial function in mediating and maintaining the higher-order organization of mTORC1 (Takahara et al., 2006; Yang et al., 2016; Yip et al., 2010). Mitotic phosphorylation of raptor may therefore regulate binding to other mTORC1 subunits and/or mTORC1 dimerization. To test this hypothesis, we expressed flag-tagged and myc-tagged mTOR in asynchronous and mitotic HeLa cells followed by Flag-IP. Flag-mTOR pulled down much less total and phosphorylated raptor from mitotic HeLa lysates as compared to asynchronous lysates (Figure 3A). Additionally, less myc-mTOR was pulled down with flag-mTOR, suggesting that raptor plays a role in dimerization of mTORC1 in mitosis. A comparable reduction in raptor in the immunoprecipitated material was observed when the antibody against mTORC1-specific component PRAS40 was used (Ramirez-Valle et al., 2010; Sancak et al., 2007) (Figure 3B). We further examined mTORC1 dimerization during mitosis by co-transfecting Flag-tagged and myc-tagged raptor. Whereas flag-tagged raptor was able to efficiently IP co-transfected myc-tagged raptor from asynchronous cell extracts, it was unable to do so from mitotic extracts (Figure 3C). Interestingly, immunoprecipitated raptor appeared to be predominantly in the unphosphorylated form, further suggesting that raptor phosphorylation affects the integrity of mTORC1 dimers. To further confirm these observations, the mTORC1 complexes from asynchronous and mitotic HeLa cell extracts were analyzed using size-exclusion chromatography (Figure 3D). mTOR eluted in two major peaks, centered at fractions 3 and 8 both in asynchronous and mitotic extracts. Based on the elution position of standards, we estimated that the peak at fraction 8 corresponds to mTORC1 dimers, whereas the peak corresponding to higher MW complexes is likely representative of mTOR multimers (Wang et al., 2006). In asynchronous cell extracts raptor co-eluted with mTOR (Figure 3D, centered on fraction 8). Interestingly, in mitotic extracts, raptor predominantly eluted at different fractions than mTOR corresponding to lower molecular weights (fractions 9 and 10). PRAS40 showed no difference in elution pattern between asynchronous and mitotic extracts. Taken together, these data show that mitotic phosphorylation of raptor affects dimerization of the mTORC1 complex.

**Figure 3:**
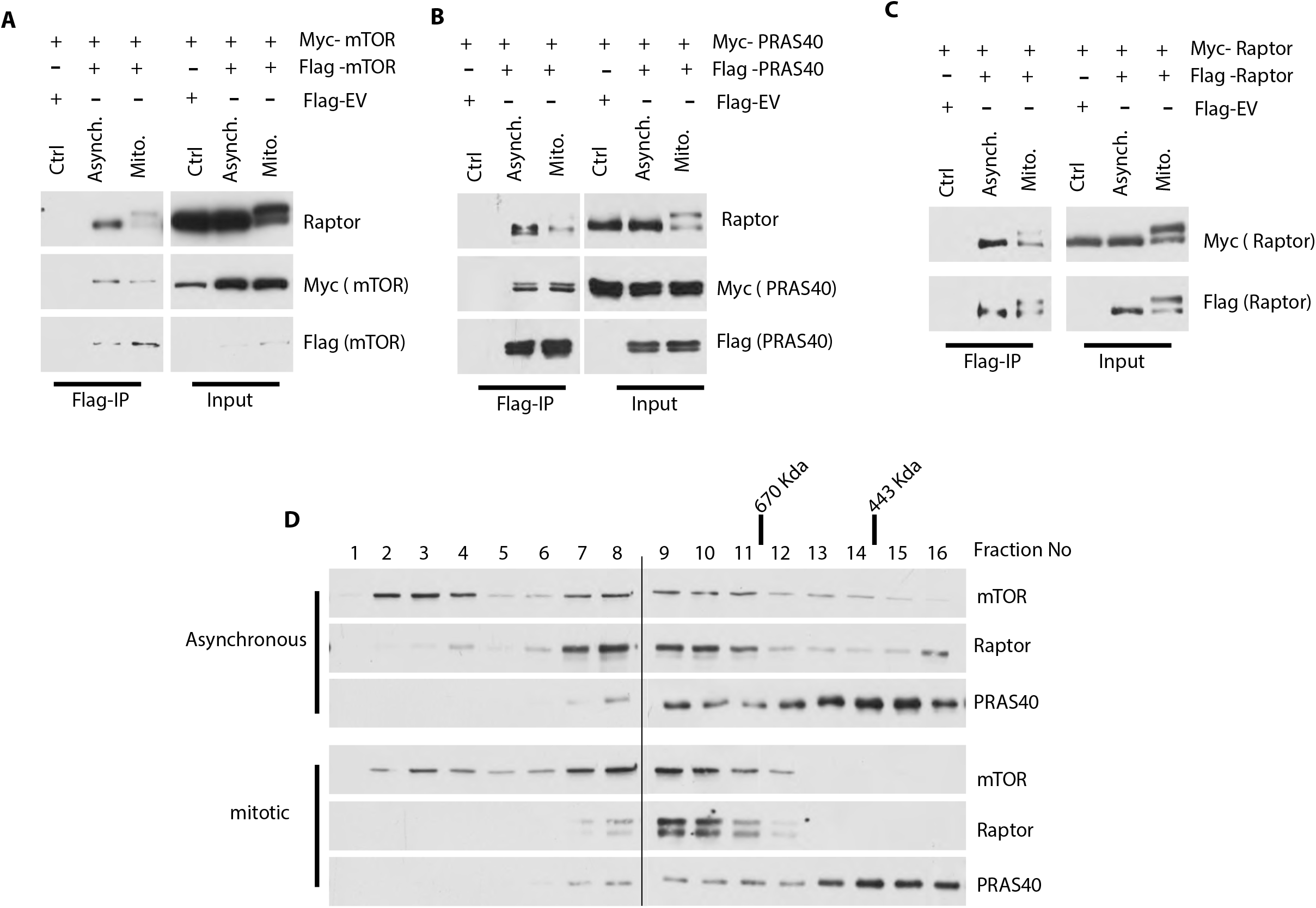
**A, B, and C-** HeLa cells were co-transfected with Flag and Myc-tagged mTOR (A), Flag and Myc-tagged PRAS40 (B), and Flag and Myc-tagged Raptor (C). Cells were synchronized as in Figure 1, cell lysates were prepared and immune-precipitation (IP) was performed using anti-Flag antibody followed by Immunoblot analysis. **D.** HeLa cells were synchronized as in A. Lysates from asynchronous (Asyn), or Thym/Noc synchronized (mito) were fractionated on a Superose 6 HR 10/30 column. Fractions were analyzed by immunoblotting with the indicated antibodies. The elution positions of molecular mass markers are shown on the top.

### Mutation of raptor phosphorylation sites activates the mTORC1 kinase in mitosis

Since we observed that raptor phosphorylation appears to negatively regulate mTORC1 in mitosis, we next identified phosphorylation sites on raptor that are responsible for this effect. Previous studies have shown that at least 9 serine and threonine residues—that cluster to two regions located between the HEAT domain and the WD40-domain of raptor—are phosphorylated in mitosis including one of the AMPK phosphorylation sites (Ser722) (Figure 4A) (Ramirez-Valle et al., 2010). Mutation of this AMPK phosphorylation site (plus another AMPK phosphorylation site, Ser792) together with other reported sites (Ser696, Thr706), however, did not exert a major effect on the electrophoretic mobility of raptor or mTORC1 activity in mitosis [Supplemental Fig. 4A]. We therefore systematically mutated all of the phosphorylation sites previously reported to occur in mitosis. These include a cluster of sites upstream (Ser696, Thr706, Ser711) and downstream (Ser855, Ser859, Ser863) of the central HEAT domain (Figure 4A). The first cluster of sites were mutated to alanine to derive the mutant raptor 3A. Similarly, the second cluster was mutated to generate the mutant raptor 3A*. The two clusters were also mutated together resulting in raptor-6A. HeLa cells were transfected with WT or mutant raptor myc-tagged cDNAs and analyzed by western blot in both mitosis and asynchronous cells. The raptor 6A mutant exhibited an attenuated mitotic electrophoretic mobility shift as compared to WT raptor, which was accompanied with a modest increase in P-Thr389 S6K (Figure 4B compare lane 2 to lane 8). Strikingly, mutation of two additional residues (Ser771 and Ser877) to generate the raptor 8A mutant completely abolished the mitotic shift of raptor and dramatically increased mitotic mTORC1 activity as monitored by S6K phosphorylation as compared to WT (Figure 4B compare lane 10 to lane 16). To confirm that these effects are mediated via the mTORC1 complex, we repeated the experiments in the presence rapamycin, which selectively inhibits mTORC1, but not mTORC2. Rapamycin treatment was able to dramatically decrease raptor 8A-induced mTORC1 reactivation in mitosis (Figure 4C). Significantly, WT or raptor 8A mutant exerted comparable effects on S6K phosphorylation in asynchronous cells, therefore suggesting that the effects of raptor 8A mutant on mTORC1/S6K axis are mitosis-specific, (Figure 4C, compare lane 1 and 3). Since we observed that mitotic phosphorylation of raptor negatively affected the formation of the active dimeric mTORC1, we examined the ability of the raptor 8A mutant to form active complex. Flag-tagged mTOR and EGFP-tagged Raptor (WT or 8A) were co-transfected in HeLa cells and flag IP was performed from asynchronous and thymidine/nocodazole arrested cells. As expected, flag-mTOR pulled down much less wild-type EGFP-Raptor from mitotic cells, confirming the negative effect of raptor phosphorylation on the assembly of mTORC1 (Figure 4D). (Figure 4D). In contrast, the EGFP-raptor 8A mutant remained associated with mTOR in mitosis and caused robust phosphorylation of S6K (Figure 4D). The effects of the raptor 8A mutant extended down to increased phosphorylation of ribosomal protein S6, which is a downstream target of the S6Ks. Taken together, these results show that phosphorylation of raptor at several sites flanking the central HEAT domain render the protein refractory to negative signals during mitosis.

**Figure 4:**
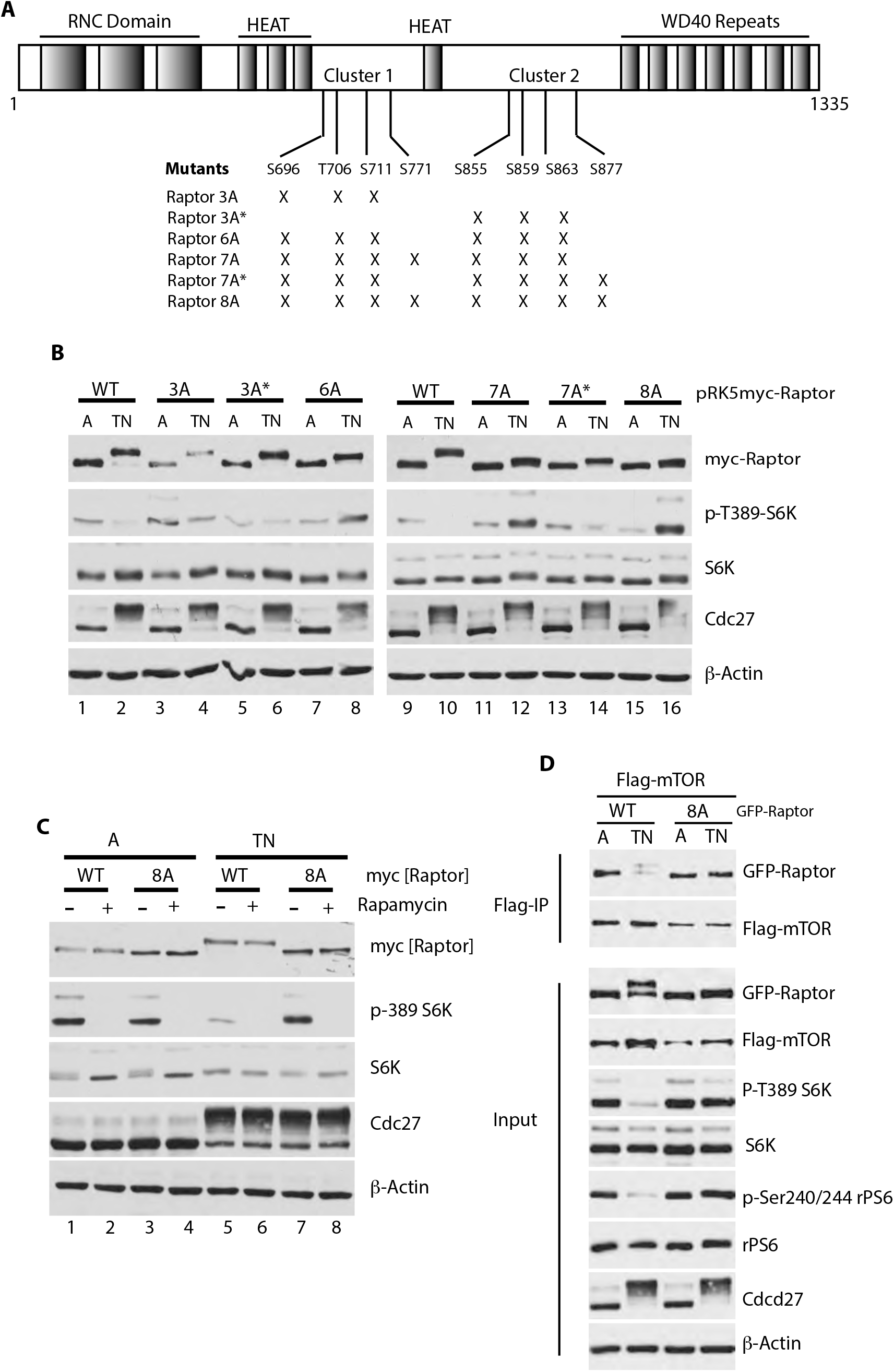
**A.** Diagram of Raptor’s protein sequence showing the mitosis-specific phosphorylation sites. **B.** Immunoblot analysis of HeLa cells that were transfected with different myc-raptor’s point mutants and were either left unsynchronized (A)or synchronized with Thymidine/Nocodazole protocol (TN). Floating mitotic cells were collected by shake-off. **C-** HeLa cells were transfected with either Raptor WT or 8A mutant, synchronized as in B, and then treated with 200nM Rapamycin for 3 hours to inhibit the mTORC1 activity. **D.** HeLa cells were co-transfected with Flag mTOR and either EGFP-Raptor WT or 8A mutant and synchronized as in B. Cell lysates were prepared and immune-precipitation (IP) was performed using anti-Flag antibody followed by Immunoblot analysis.

### Active mTORC1 promotes survival in response to mitotic poisons

Several studies have shown that under prolonged mitotic arrest global rates of protein synthesis are strongly reduced (Dobrikov et al., 2014; Pyronnet et al., 2001; Wilker et al., 2007). Since mTORC1 increases the rates of mRNA translation, we next assessed the effect of raptor-8A mutant overexpression on the overall protein synthesis in HeLa cells during mitotic arrest. In mitotic HeLa cells, raptor 8A increased global protein synthesis by approximately 15% compared to WT (Figure 5A). In order to determine whether the raptor 8A mutant affects normal mitotic progression, WT myc-raptor and the 8A mutant were transfected into HeLa cell and synchronized with thym/noc, and then released into fresh media (Figure 5B). As expected, compared to WT, myc-raptor 8A showed a sustained signal for p-Thr389 S6K, starting in mitosis which was maintained throughout exit and entrance into G1. The timing for cyclin-B1 degradation (marking mitotic exit) showed no major differences between WT and 8A myc-raptor expressing cells. Similarly, time lapse microscopy of cells transfected with EGFP-raptor WT or EGFP-raptor 8A displayed only a slight increase in the average time taken for the mutant expressing cells to transit through one complete cell cycle (Figure 5C).

**Figure 5:**
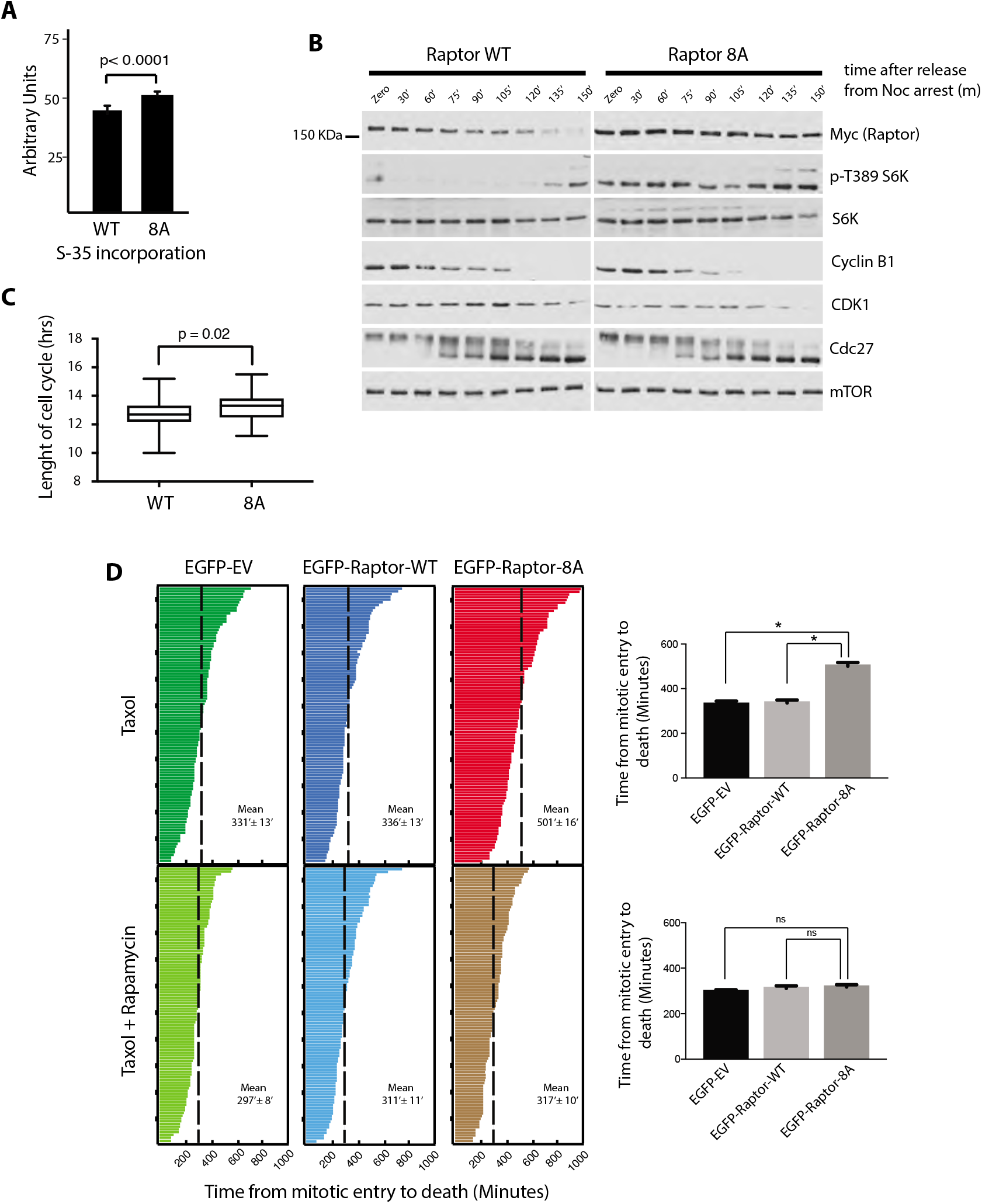
**A.** HeLa cells were transfected in triplicates with either pRK5-myc raptor WT or 8A mutant and synchronized with Thym/Noc protocol then labeled with [S^35^] methionine and the specific activity of incorporation into equal amounts of protein was determined by trichloroacetic acid (TCA) precipitation and scintillation counting. **B.** Immunoblot analysis of HeLa cells that were transfected with pRK5-myc raptor WT or 8A synchronized with Thym/Noc protocol. Nocodazole arrested cells were collected and released into fresh media. Degradation of cyclin B1, and phosphorylation of cdc27 are used as marker for exit from mitosis. **C.** The length of one whole cell cycle (between two mitosis) was measured for 50 cells (transfected with, EGFP raptor WT, or 8A) using time-lapse microscopy. **D.** HeLa cells were transfected with either EGFP-empty vector or Raptor WT or 8A mutant. Twenty-four hours following transfection, cells were synchronized by RO3306. After 20 hours cells were washed twice with PBS and released into fresh media containing either 100 nM Taxol (Top panel) or 100 nM Taxol + 200nM Rapamycin, and time-lapse imaging started. The length of time spent by 100 cells from mitotic entry until death was plotted.

Since mTORC1 signaling is known to promote survival by stimulating translation and inhibiting autophagy, we hypothesized that the raptor 8A mutant may affect cell fate during mitotic arrest. To this end, HeLa cells were transfected as above, synchronized in G2/M using RO3306, and then released into fresh media containing 100nM taxol with or without 200nM rapamycin. The fates of GFP positive cells were monitored using time-lapse imaging. In taxol-treated cells the average time cells remained arrested in mitosis before death was significantly higher in cells transfected with EGFP-raptor 8A relative to empty vector or EGFP-raptor WT (Figure 5D). This effect was mTORC1 dependent since addition of rapamycin reversed the pro-survival effects induced by raptor-8A mutant (Figure 5D). These data show that preventing mTORC1 inhibition during prolonged mitotic arrest enhances cell survival.

### PDCD4 mediates survival downstream of active mTORC1 in response to Taxol

To further investigate the mechanism(s) by which the raptor 8A mutant attenuates cell death during mitotic arrest, HeLa cells transfected with either empty EGPG vector, EGFP-raptor WT, or EGFP-raptor 8A were maintained under asynchronous or taxol-arrested mitotic conditions. Analysis of known mTORC1-regulated proteins that control cell survival revealed that Bcl-xL slightly increased and PDCD4 dramatically decreased in EGFP-raptor 8A mutant transfected as compared to EGFP-raptor WT transfected or control cells (Figure 6A, compare lanes 5 and 6). PDCD4 is a known tumor suppressor (Lankat-Buttgereit and Goke, 2003) that functions as an inhibitor of eukaryotic translation initiation factor 4A (eIF4A) (Yang et al., 2003). PDCD4 is phosphorylated via the mTORC1/S6K axis and its proteasomal degradation is mediated by SCF^βTrCP1^ E3 Ligase (Dorrello et al., 2006). To confirm the correlation between PDCD4 and active mTORC1 in mitosis we depleted endogenous PDCD4 using two different siRNAs. Cells were synchronized as in figure 5D, released into 100nM taxol and analyzed by time-lapse imaging. Strikingly, depletion of PDCD4 phenocopied the effects of expressing the raptor-8A mutant, resulting in a significant prolongation in mitotic survival during taxol treatment (Figure 6B). Since PDCD4 is known to inhibit the activity of eIF4A helicase activity, we hypothesized that eIF4A may be involved in mediating the pro-survival activity of raptor 8A. To address this question, we repeated the same experiment as in figure 5D in the presence of hippuristanol, a selective eIF4A inhibitor. Similar to rapamycin, hippuristanol reversed the effects of raptor 8A mutant on survival during prolonged mitotic arrest (Figure 5C). These data suggest that the pro-survival effects of raptor 8A during mitotic arrest are mediated by the downregulation of PDCD4 expression and subsequent increased activity of eIF4A.

**Figure 6:**
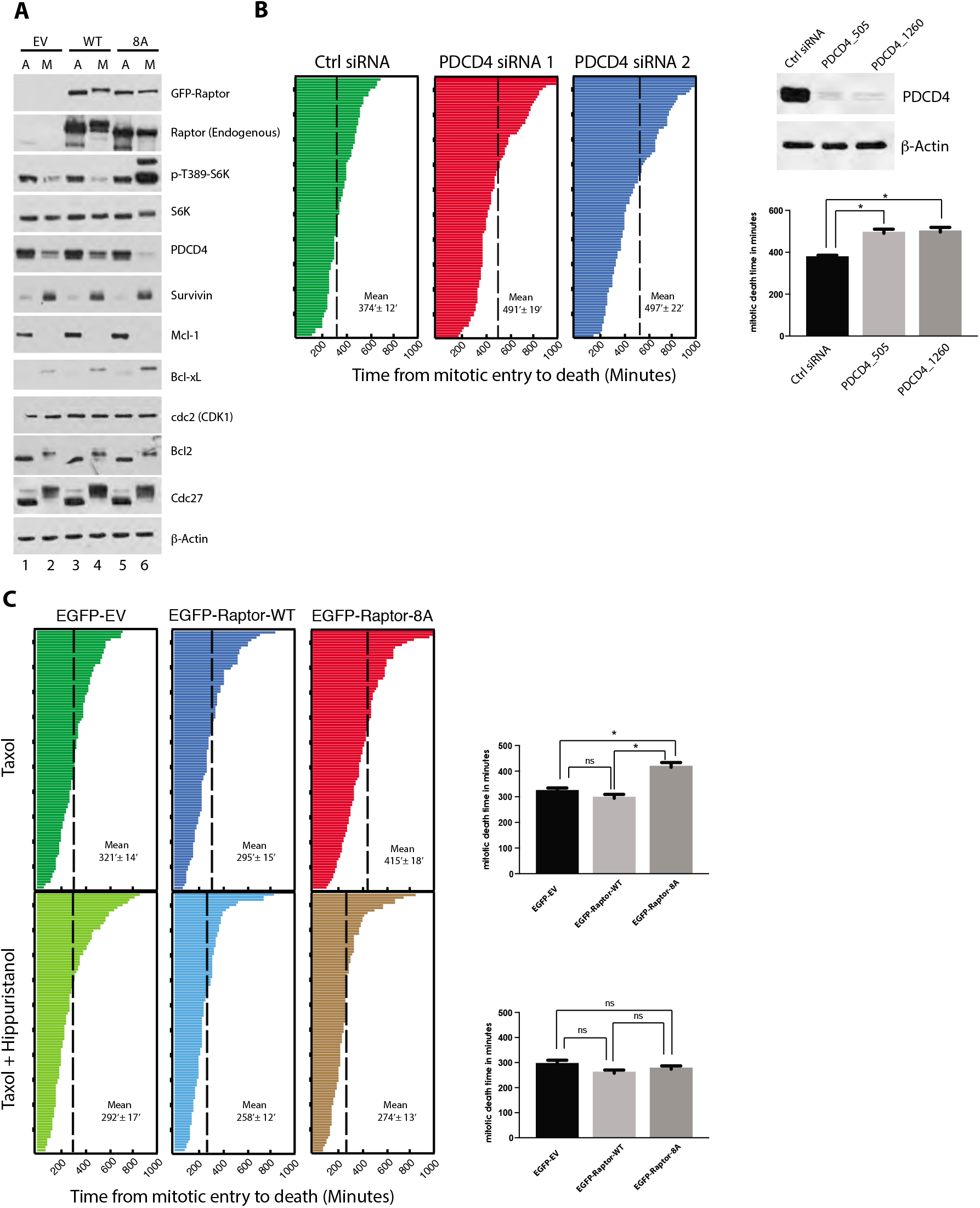
**A.** Immunoblot analysis of HeLa cells transfected with either EGFP-empty vector or Raptor WT or 8A mutant. Twenty-four hours following transfection, cells were synchronized using Thym/Noc protocol. Cells harvested after 15 hours and lysates immunoblotted for the indicated antibodies. **B.** HeLa cells were transfected overnight with 50 nM siRNAs as indicated. Twenty-four hours following transfection, cells were synchronized by RO3306. After 20 hours cells were washed twice with PBS and released into fresh media containing 100nM Taxol, and time-lapse imaging started. The length of time spent by 100 cells from mitotic entry until death was plotted. The knockdown was confirmed by immunoblotting using PDCD4 antibody. **C.** HeLa cells were transfected with either EGFP-empty vector or Raptor WT or 8A mutant. Twenty-four hours following transfection, cells were synchronized by RO3306. After 20 hours cells were washed twice with PBS and released into fresh media containing either 100 nM Taxol (Top panel) or 100 nM Taxol + 200nM Hippuristanol, and time-lapse imaging started. The length of time spent by 100 cells from mitotic entry until death was plotted. The mean time for cell death is compared to the right for EGFP-EV, raptor WT, and raptor 8A in a bar graph using student t-test (two-tailed, unpaired).

### The role of mTOR/S6K/PDCD4/eIF4A axis in regulating death vs. slippage decisions upon prolonged mitotic arrest

The current model for cell fate during extended mitotic arrest proposes that there is competition between death and mitotic exit with each having its own threshold (Gascoigne and Taylor, 2008). If a cell accumulates enough death signals, the death threshold is reached first, and the cell dies. Conversely, if the cells cannot maintain the mitotic state and the mitotic exit threshold is surpassed, cells exit without cell division (slippage). The observation that hippuristanol can abolish the prosurvival effects of mTORC1 during mitosis arrest prompted us to test if the mTORC1/S6K/PDCD4/eIF4A pathway is involved as a timer for the death threshold during mitotic arrest.

To test this hypothesis, HeLa cells were arrested in G2 with RO3306 and released into 100nM taxol or taxol and varying concentrations of hippuristanol. Figure 7A shows that hippuristanol acted in a dose dependent manner to reduce the time to cell death when combined with taxol. The mean time of death for a hundred cells was reduced from 448 minutes with taxol alone to only 188 minutes when taxol and 500 nM hippuristanol were combined. Whereas treatment with 100nM taxol causes HeLa cells to die during mitotic arrest, lower concentrations (10 nM) predominately favored slippage (Gascoigne and Taylor, 2008). We therefore determined if eIF4A inhibition could shift the fate of HeLa cells treated with low dose Taxol towards death. Figure 7B shows that treatment of HeLa cells with 10nM taxol resulted in most cells undergoing slippage with only 24% of cell undergoing cell death. Combined taxol/hippuristanol treatment attenuated this effect and resulted in 71% death in mitosis with only 29% slippage. To exclude potential cell line biases and generalize these findings, we extended the investigation to two other cancer cell lines: H1299, a small cell lung carcinoma, known to exhibit slippage in response to taxol, and MCF-7, an invasive breast ductal carcinoma. In response to 300 nM taxol, H1299 cells mainly underwent slippage (74%) with just 22% cell death. Adding 500nM hippuristanol increased the death to 69% and reduces slippage to just 12% (Figure 7C). Similarly, MCF7 cells displayed an increase in the percentage of death when treated with the drug combination (68%), as compared to taxol alone (42%) (Figure 7D).

**Figure 7:**
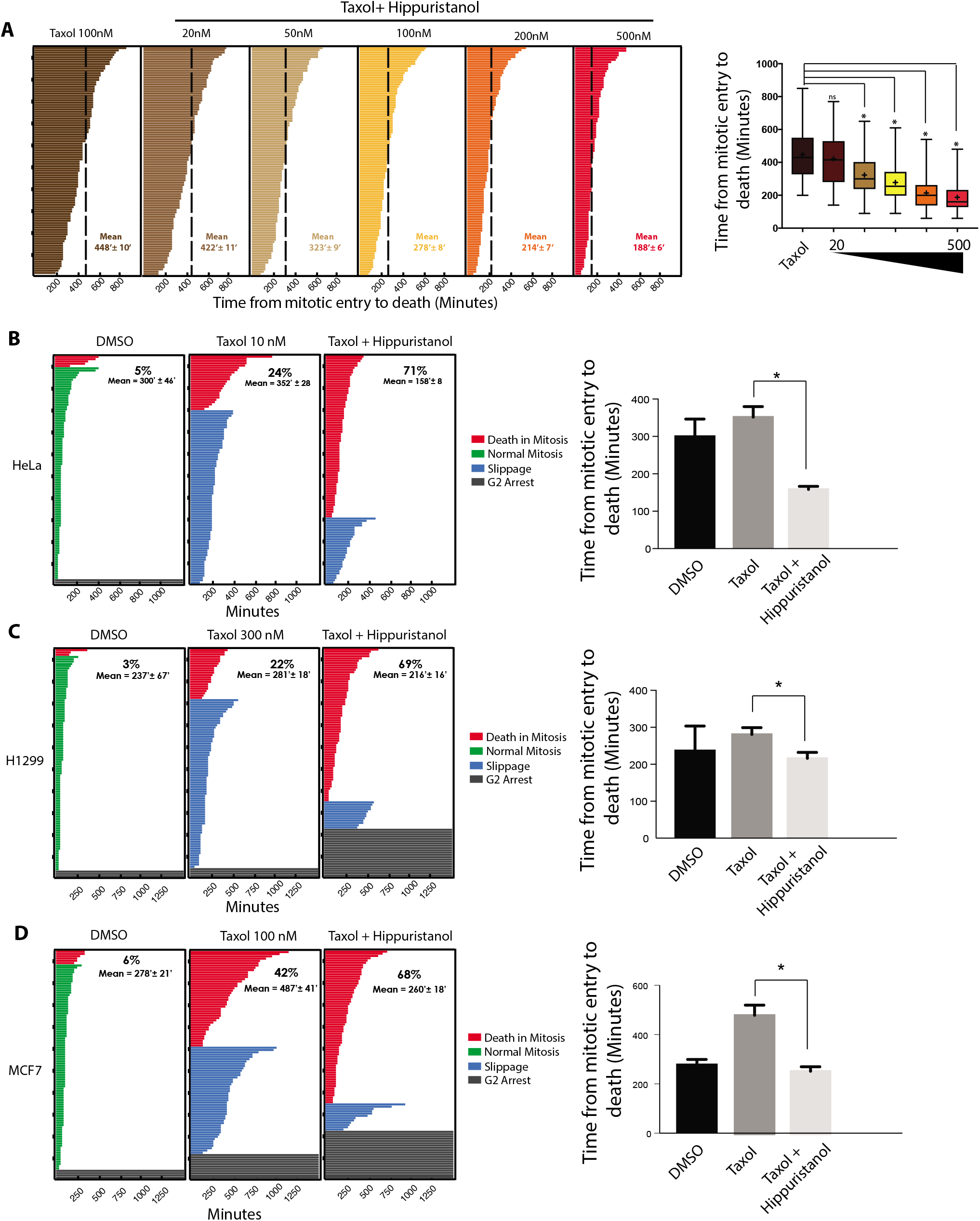
**A-** HeLa cells were synchronized by RO3306. After 20 hours cells were washed twice with PBS and released into fresh media containing either 100 nM Taxol (Top panel) or 100 nM Taxol plus increasing concentrations of Hippuristanol as indicated, and time-lapse imaging started. The length of time spent by 100 cells from mitotic entry until death was plotted (left panel). A box plot comparing the range and the mean death time for all of the treatment is shown to the left. **B.** HeLa cells were synchronized as in A but then released into either Taxol (10nM), or Taxol plus Hippuristanol (200nM), and then time-lapse analysis started as in A. **C**. H1299 cells were synchronized and treated as in B then released into either Taxol (300nM), or Taxol plus Hippuristanol (500nM), followed by time-lapse analysis. **D**. MCF-7 cell were synchronized with Thymidine for 20 hours followed by release into fresh media for three hours and then addition of the drug as indicated. Time lapse imaging was then started. The fate of 100 cells is shown for B, C, and D. The mean time for cell death is compared to the right for each cell lines in a bar graph using student t-test (two-tailed, unpaired).

Since hippuristanol appeared to enhance the anti-mitotic effects of taxol, we examined the effect of combined treatment of H1299 and MCF7 cells on cell growth. Interestingly, hippuristanol was able to reverse the resistance of H1299 to taxol and greatly enhanced the effects of even low dose (50 nM) taxol. Similar effects were observed with MCF7 cells (Supplemental 7A).

In summary, these findings confirm our observation that cells undergoing prolonged mitotic arrest require shut down the mTORC1 activity in order to maintain PDCD4 levels and eIF4A activity. We speculate that mTORC1/PDCD4/eIF4A regulation during prolonged mitotic arrest may serve as a checkpoint to prevent slippage and ensuing genomic instability.

## Discussion

Previous studies have shown that raptor is phosphorylated during prolonged mitotic arrest, but the significance of this phosphorylation has been largely unknown. In the current study we show that phosphorylation of raptor at multiple sites, plays a crucial role in deactivating the mTORC1 complex during prolonged mitotic arrest. A non-phosphorylatable raptor mutant (raptor 8A), can restore the activity of mTORC1 complex and promotes survival of mitotic cells when exposed to mitotic poisons. The effect of the raptor 8A mutant is mediated at least in part by down-regulation of PDCD4 levels and consequent increase in the activity of its downstream target, eIF4A. This mechanism of mitotic mTORC1 inactivation is similar to that observed after of energy deprivation and activation of the AMPK pathway in which AMPK-mediated phosphorylation of raptor induces 14-3-3 binding and inhibition of mTORC1 (Gwinn et al., 2008). However, a role for AMPK mitotic inhibition of mTORC1 was ruled out by the use of the raptor Ser 722/792 mutant, which retained activity in mitosis.

In general. the duration of a normal mitosis is approximately one hour, so it is unlikely that the effects of mTORC1 signaling on mRNA translation would have major effects during this timeframe. However, during a condition of aberrant mitosis in which the mitotic checkpoint is engaged for an extended period, lack of mTORC1 signaling would negatively affect mRNA translation. We observed that during prolonged mitotic arrest, not only is raptor inactivated by phosphorylation, its steady state level is also appeared to be reduced in a range of cancer and normal cell lines. We hypothesize that shut down of mTORC1 activity during extended mitotic arrest may act as a checkpoint to prevent excessive survival and slippage of cells undergoing aberrant mitosis. Slippage through mitotic arrest results instantly in genomic duplication, which is thought to be a contributing factor to the process of tumorigenesis (reviewed in (Lens and Medema, 2019)).

Translation is one of the most energy consuming processes in the cell (Buttgereit and Brand, 1995), which may also explain the reason behind inhibition of mTORC1 during prolonged mitotic arrest as a mechanism to cope with energy stress. Suppression of other core components that regulate translation including eIF4E and eIF4G have been previously reported (Bonneau and Sonenberg, 1987; Dobrikov et al., 2014). Since mTORC1 is a positive regulator of cap-dependent-translation, inhibition of mTORC1 fits into the general effect of limiting translation capacity of cells during mitosis.

Our data also show that the tumor suppressor PDCD4 level is highly sensitive to mTORC1 activity in mitosis which agrees with previous reports showing its degradation through the mTORC1/S6K axis (Dorrello et al., 2006). PDCD4 is a known inhibitor of eIF4A (Yang et al., 2003). Mitotic arrested cells need to maintain a reduced level of translation (Mena et al., 2010; Zeng et al., 2010). This residual translation may be involved in determining the fate of the cells under prolonged arrest which is determined by the competition between two networks; one regulates the buildup of apoptotic signals and the other promotes exit without cytokinesis (mitotic slippage) (Gascoigne and Taylor, 2008).

Targeting eIF4A activity in the presence of taxol resulted in cells predominantly undergoing death rather than slippage. Pharmacological inhibition of eIF4A also led to more rapid death in a dose-dependent manner which may explain how cells time induction of death during mitotic arrest. For example Bcl-xL, an anti-apoptotic protein have been shown to be an “eIF4A-sensitive” target and is selectively up-regulated when eIF4A activity is increased (Gandin et al., 2016; Li et al., 2003). Lowering eIF4A activity would therefore be expected to reduce levels of Bcl-xL and other anti-apoptotic proteins.

Although drugs targeting the microtubules like taxol and vincristine are front line therapies for several cancers, their toxicity represents a large limitation. Our results show that it is possible to obtain the same sensitivity of cancer cells to these drugs but using a much lower dose. Combining taxol with other drugs targeting the translation machinery such as eIF4A inhibitors may have clinical application in cancers that are otherwise resistant to taxol or other anti-mitotics.

## Experimental Procedures

### Cell lines and treatments

Cells were maintained in Dulbecco’s modified Eagle medium (Wisent Inc., QC, Canada) supplemented with 10% fetal bovine serum (HyClone; Thermo Scientific) and 0.1% gentamicin (Wisent Inc., QC, Canada). Cells synchronized in mitosis were obtained by 20 hours treatment with 2.5 mM thymidine (Sigma), released for 4 hours, then treated with 100 ng/ml nocodazole (Sigma). RO3306 synchronization was performed by treating HeLa cells for 20 hours with RO3306 (Enzo). proTAME (Boston Biochem) was used at 10 to 20 μM. Taxol (Sigma) was used at 100nM unless otherwise indicated. MG132 (Sigma) was used for 4-hour treatments. Other drugs that used included: Chloroquine, Baffylomycin, ammonium chloride, rapamycin (200nM) and Hippuristanol (Dr. Jerry Pelletier). Treatment of cell extracts with λ phosphatase (New England BioLabs) was performed by adding 50 μg of total protein extract to the 1×phosphatase buffer (supplied by manufacturer) supplemented with 2 mM MnCh and incubated with 10 U of □ phosphatase at 30°C for 30 minutes. The reactions were stopped by the addition of 1x Laemmli sample buffer.

### Plasmids and Cloning

pRK5-Myc-empty vector was purchased from Clontech. The following vectors were purchased from Addgene: Raptor wt (Plasmid #1859), myc-Raptor S722A/S792A (Plasmid #18118), pcDNA3-Flag mTOR wt (Plasmid #26603), myc-mTOR (Plasmid #1861), pRK5 Flag PRAS40 (Plasmid #14950), pRK5-myc-PRAS40 (Plasmid #15476). Raptor point mutations were generated using site-directed mutagenesis. To generate the EGFP-Raptor fusion protein, Raptor was sub-cloned into pEGFPC1 (Clontech). All constructs were verified by restriction digest analysis and DNA sequencing.

### Extract Preparation and Immuno-precipitation

Cell extracts were prepared by lysing cells in Lysis Buffer (50 mM Tris pH 7.5, 150 mM NaCl, 0.5% NP40, 1 mM EDTA, 1 mM Na_3_VO_4_, 50 mM NaF, 10 mM β-glycerophosphate, 1 mM PMSF and protease inhibitor (Roche)), followed by protein quantification via Bradford assay (Bio-rad). For Flag-mTOR, Flag PRAS40 or Flag-Raptor immunoprecipitations, transfected HeLa or Hela cells were lysed in 1 ml IP Lysis buffer (40 mM HEPES pH7.4, 100mM NaCl, 0.3% CHAPS, 2mM EDTA, 10 mM Sodium pyrophosphate and protease inhibitor (Roche)) per 10^7^ cells on ice for 20 min. Cell debris was pelleted by centrifugation at maximum speed for 15 min at 4°C. The supernatant was then incubated with 30 μl anti-Flag (Sigma) for 2 h at 4°C. The beads were then pelleted and washed 4 times with the same buffer. Following the washes, the beads were pelleted and resuspended in 1× Laemmli sample buffer, boiled for 5 min, and stored at −20°C until further use.

### Antibodies and Blotting

Mouse monoclonal antibodies to the following proteins were purchased from the indicated manufacturers and used for immunoblotting according to standard protocols: cyclin B1, cyclin A2 (Santa Cruz Biotechnology), Cdc27 (BD Biosciences), EGFP (Clontech), anti-Flag M2, (Sigma), anti-myc (Bioshop), Cdk1, ribosomal S6(Cell Signaling), Bcl2 (Zymed). The following rabbit polyclonal antibodies were purchased from the indicated manufacturers: Raptor (Millipore), β-actin (Sigma),p70-S6K(SC), phospho-4EBP1/2Thr37/46, Phospho-S6K-Thr389, Geminin, P-Ser2448 mTOR, eIF4B, P-Ser240/244 rPS6(Cell Signalling). Rabbit monoclonal antibodies were from Cell Signaling (mTOR,4EBP1/2, TSC2, PRAS40 and PDCD4). Western blotting was performed using standard protocols for SDS-PAGE and wet transfer for at least 24 hours at 30V onto nitrocellulose membranes (Bio-Rad). Conditions for Western blots were the use of 5% nonfat dry milk in TBS-T (50 mM Tris [pH 7.5], 150 mM NaCl, 0.5% Tween 20). The bands were visualized by enhanced chemiluminescence (Western Lightning [PerkinElmer] or SuperSignal™ West Femto [ThermoFisher Scientific]) and exposure on film.

### Gel Filtration Chromatography

About 4.0 x10^7^ HeLa cells were lysed with 0.3 ml of lysis buffer on ice for 20 min. After centrifugation at 13,000 g for 10 min, the supernatant was passed through a 0.22-m filter (Millipore Corp., Bedford, MA). About 1.5–2.0 mg of proteins in 0.3-ml volume were loaded onto a Superose 6 HR10/30 column (GE HealthCare) pre-equilibrated with lysis buffer. The proteins were eluted at 0.2 ml/min, and 0.5-ml fractions were collected. Sixteen microliters of each were then analyzed by immunoblotting with the indicated antibodies.

### siRNA transfection

150,000 HeLa cells were seeded in 6-well plate and then transfected overnight using 5μl Lipofectamine 2000 (Invitrogen-Life Technologies) per well and 50 nM siRNA in Opti-MEM reduced-serum medium (Invitrogen-Life Technologies). All siRNAs were purchased from Sigma. The cells were allowed to recover and then treated as indicated. PDCD4_505 5’CAC CAA UCA UAC AGG AAU A dTdT3’ (Wang et al., 2015) PDCD4_1260-5’CAU UCA UAC UCU GUG CUG G dTdT3’ (Bitomsky et al., 2008) Non-silencing 5’-AAT TCT CCG AAC GTG TCA CGT dTdT-3’ (Qiagen)

### [^35^S]methionine-cysteine pulse-chase

HeLa cells were transfected and synchronized as above. 15 hours post Nocodazole floating cell were collected by shake off, cells pelleted and resuspended in DMEM without methionine and cysteine and supplemented with 10% dialyzed serum (both from Gibco) for 2 h and 100ng/ml nocodazole and replated. Cells were then labeled with a 10 μCi/ml mixture of [^35^S]methionine-cysteine (Amersham) for 10 min. Cell lysis was performed as previously described, and 10μl of supernatant was precipitated by trichloroacetic acid (TCA) on a filter paper. Filter papers were soaked in scintillation fluid, and radioactivity was measured using a β-scintillation counter (Gandin et al., 2013).

### Real Time Quantitative PCR analysis

Total RNAs were extracted from HeLa cells using Trizole protocol.1ug RNA was reverse-transcribed to cDNA (QuantiTect Reverse Transcription Kit, Qiagen). Primers used are: Raptor-Fwd 5’GCCTGCTGTACATAGTGAAGCT 3’, Raptor-Rev 5’ TGGATGCTGGTGCTCAGTGGG3’, 18S-Frd 5’GTAACCCGTTGAACCCCATT 3’ and 18S-Rev 5’ CCATCCAATCGGTAGTAGCG 3’. QRTPCR analysis was performed on the Eppendorf realplex using the QuantiTect SYBR Green (Qiagen). Gene expression analysis was determined using the delta CT method and normalized to 18S.

### Time-Lapse Microscopy

100,000 HeLa cells (125,000 for H1299 and MCF7) were seeded on 6-well plates and then either transfected or treated with drugs as indicated. Synchronization was performed as described above. Following the addition of Taxol or Hippuristanol, the cells were placed in an incubation chamber on the microscope to maintain temperature and CO_2_ levels. Images were taken every 10 minutes at 10X total magnification. For analysis, 100 cells were followed for each condition and the outcome of mitosis recorded

---

**Supplementary figure 1:**
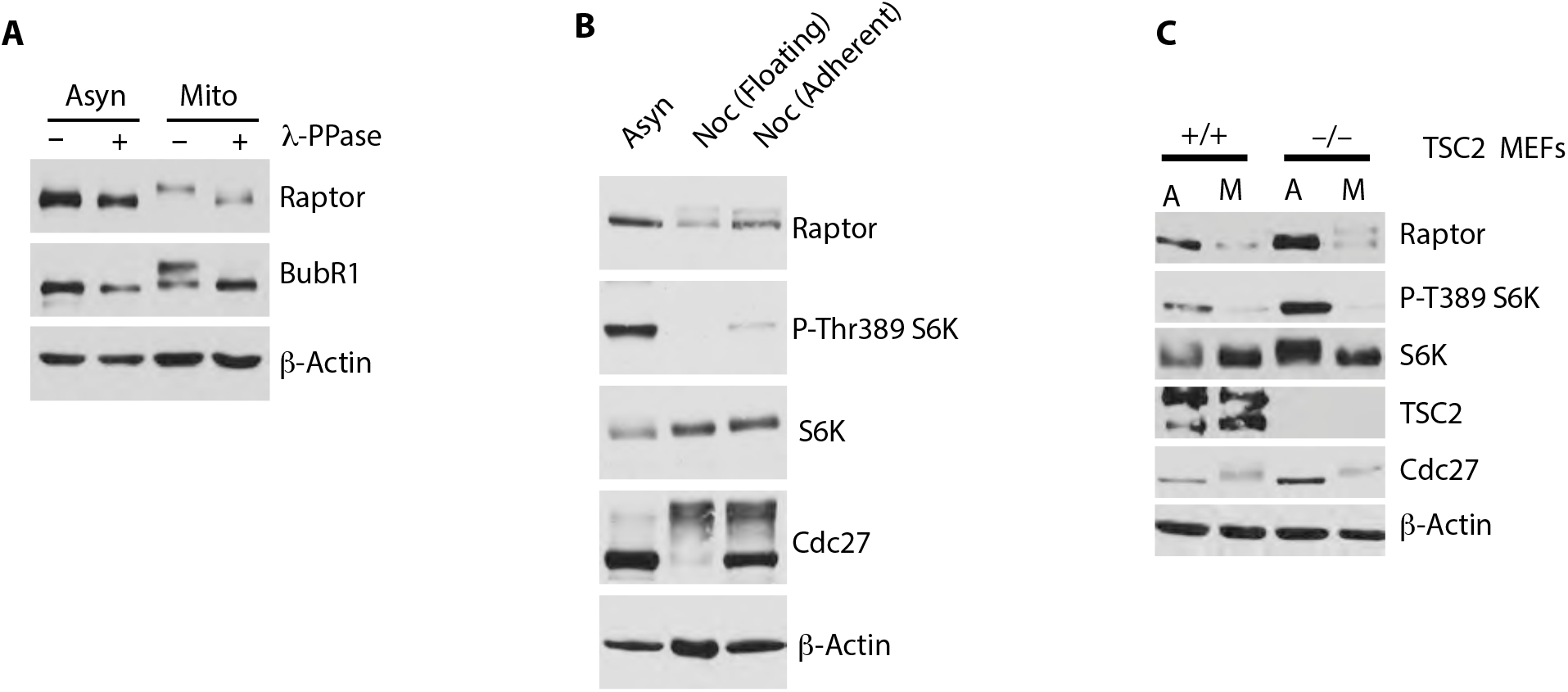
**A.** Immunoblot analysis of endogenous Raptor in whole cell extracts treated with λ phosphatase (+) or control (-) prepared from asynchronous or Thymidine/ Nocodazole mitotic extracts. BubR1 phosphorylation was used as a control for the phosphatase. **B.** Immunoblot of HaCat cells that were either left unsynchronized (Asyn) or synchronized using Thymidine-Nocodazole protocol. 15 hours post-Nocodazole mitotic cells were collected by shake-off (Floating) or scraped from their plate (Adherent). **C.** WT and TSC2 Null MEFS were synchronized as in A except that they were released in 100nM Taxol and harvested by shake-off 60’ after release.

**Supplementary figure 2.**
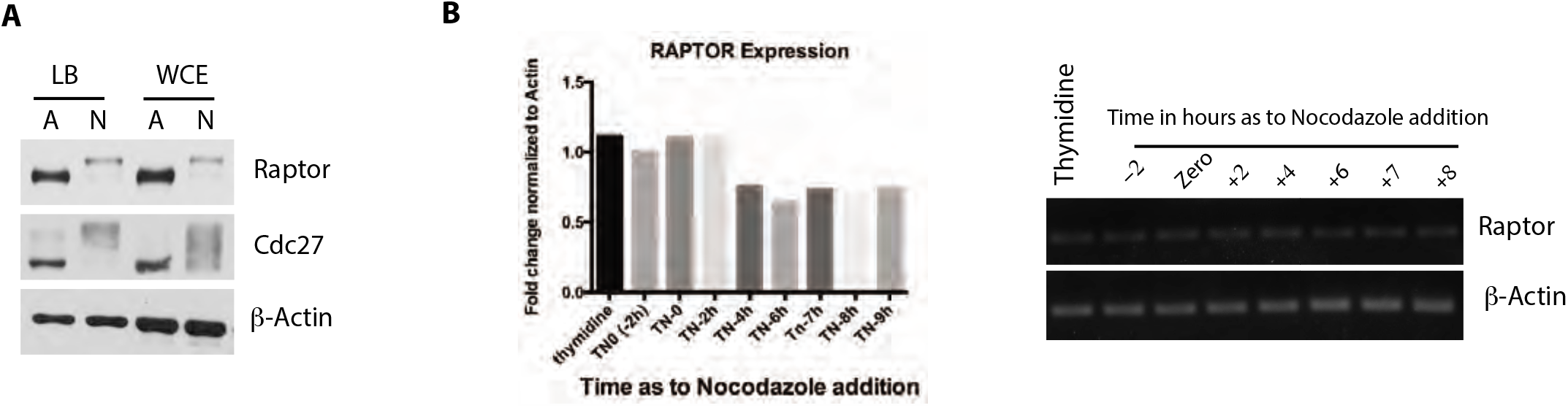
**A.** Immunoblot of HeLa cells that were either left unsynchronized (Asyn) or synchronized using Thymidine-Nocodazole protocol. 15 hours post-Nocodazole, mitotic cells were collected by shake-off and asynchronous cells by scraping. Cell pellet was either lysed in lysis buffer or whole cell extract was prepared in Laemmli buffer. **B.** Expression analysis of raptor’s mRNA level in HeLa cells under Thymidine or released from thymidine into Nocodazole and Harvested at different time points.

**Supplementary figure 4.**
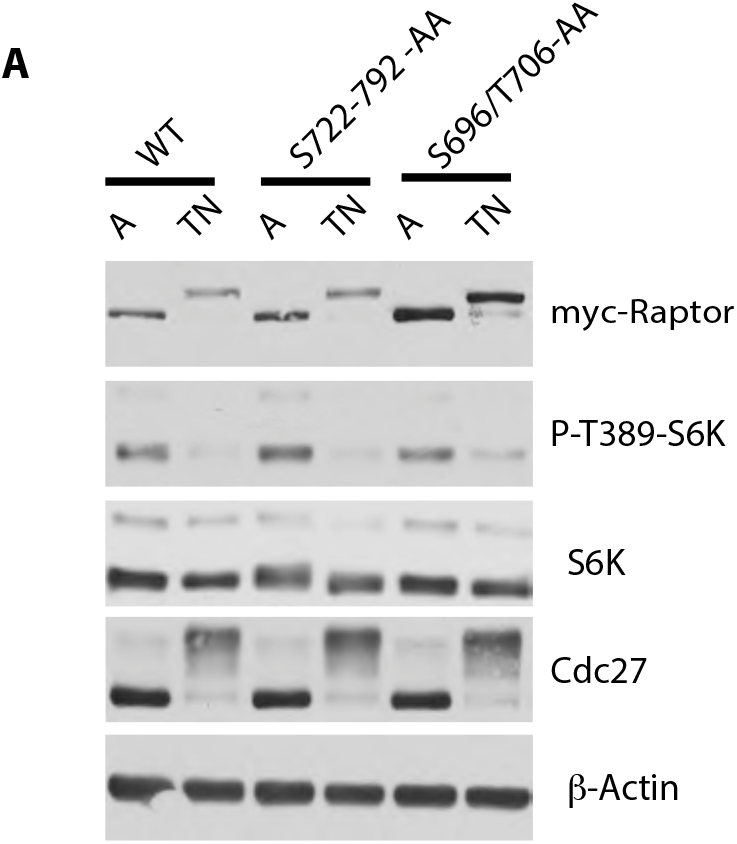
**A.** Immunoblot analysis of HeLa cells that were transfected with different myc-raptor’s point mutants and were either left unsynchronized (A)or synchronized with Thymidine/Nocodazole protocol (TN).

**Supplementary figure 7.**
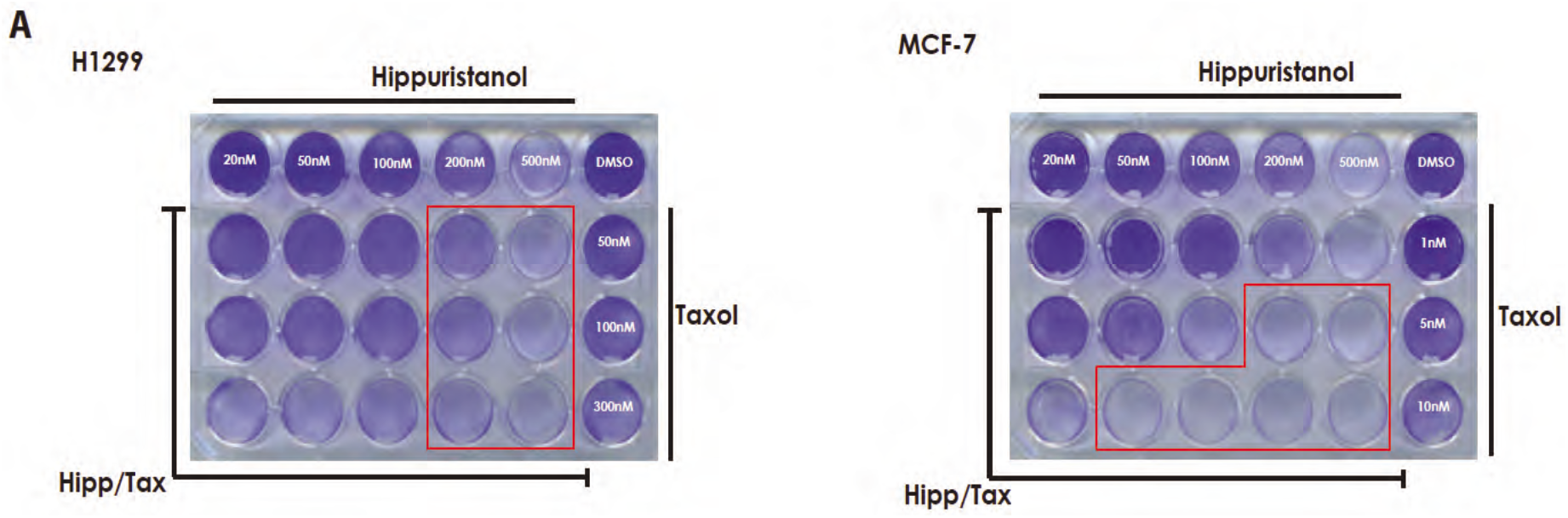
**A.** Crystal violet staining of H1299 cells (left) or MCF7 (right) treated with different concentrations of hippuristanol, taxol, or combination of both. No synchronization protocol was applied, and cells were stained 72 hours post treatment.

